# Beyond gait speed: a multidimensional motor signature of Motoric Cognitive Risk syndrome identified through domain-specific anomaly detection

**DOI:** 10.64898/2026.04.22.716304

**Authors:** Baptiste Perthuy, Hugues Vinzant, Clément Brifault, Vincent Cabibel, Rémi Laillier, Pierre Denise, Nicolas Lefèvre, Alexandre Dalibot, Nick Stergiou, Fabien Cignetti, Leslie M. Decker

## Abstract

Motoric Cognitive Risk (MCR) syndrome, defined by subjective cognitive complaints and slow gait speed, identifies older adults at increased risk of major neurocognitive disorders (NCDs). Yet, gait speed reflects a composite output shaped by heterogeneous neuromusculoskeletal and cognitive processes, limiting its clinical specificity. This study aimed to refine the motor signature of MCR by quantifying domain-specific gait deviations relative to a normative reference cohort using an anomaly detection approach.

Ninety-seven adults (≥ 55 years) completed two 3-minute treadmill walking bouts at their preferred speed. Participants were categorized into three groups: older adults with MCR (n = 20), healthy older adults with slow gait (sHOA; n = 20) matched to MCR for age and gait speed, and healthy older adults (HOA; n = 57). Linear spatiotemporal and nonlinear trunk acceleration-derived variables were organized into ten functional gait domains, conceptually grouped into gait pattern (pace, rhythm, phases, postural control, symmetry), fluctuation amplitude (variability), and temporal structure of fluctuations (regulation, signal complexity, divergence of movement trajectories, and attractor complexity). For each domain, a Gaussian mixture model trained on HOA data defined a normative reference space, from which individual anomaly scores quantified deviations across groups.

Both sHOA and MCR showed higher deviations in gait pattern domains (pace and phases) than HOA, consistent with their slower gait speed. Only MCR exhibited additional deviations in domains related to fluctuation amplitude and temporal structure, reflected by increased step-to-step variability and trunk acceleration fluctuations that were more divergent, more predictable, and less complex.

These findings reveal a multidimensional motor signature of MCR. Domain-specific anomaly scores may provide individualized, clinically interpretable biomarkers to support early detection and monitoring of older adults at increased risk of major NCDs.

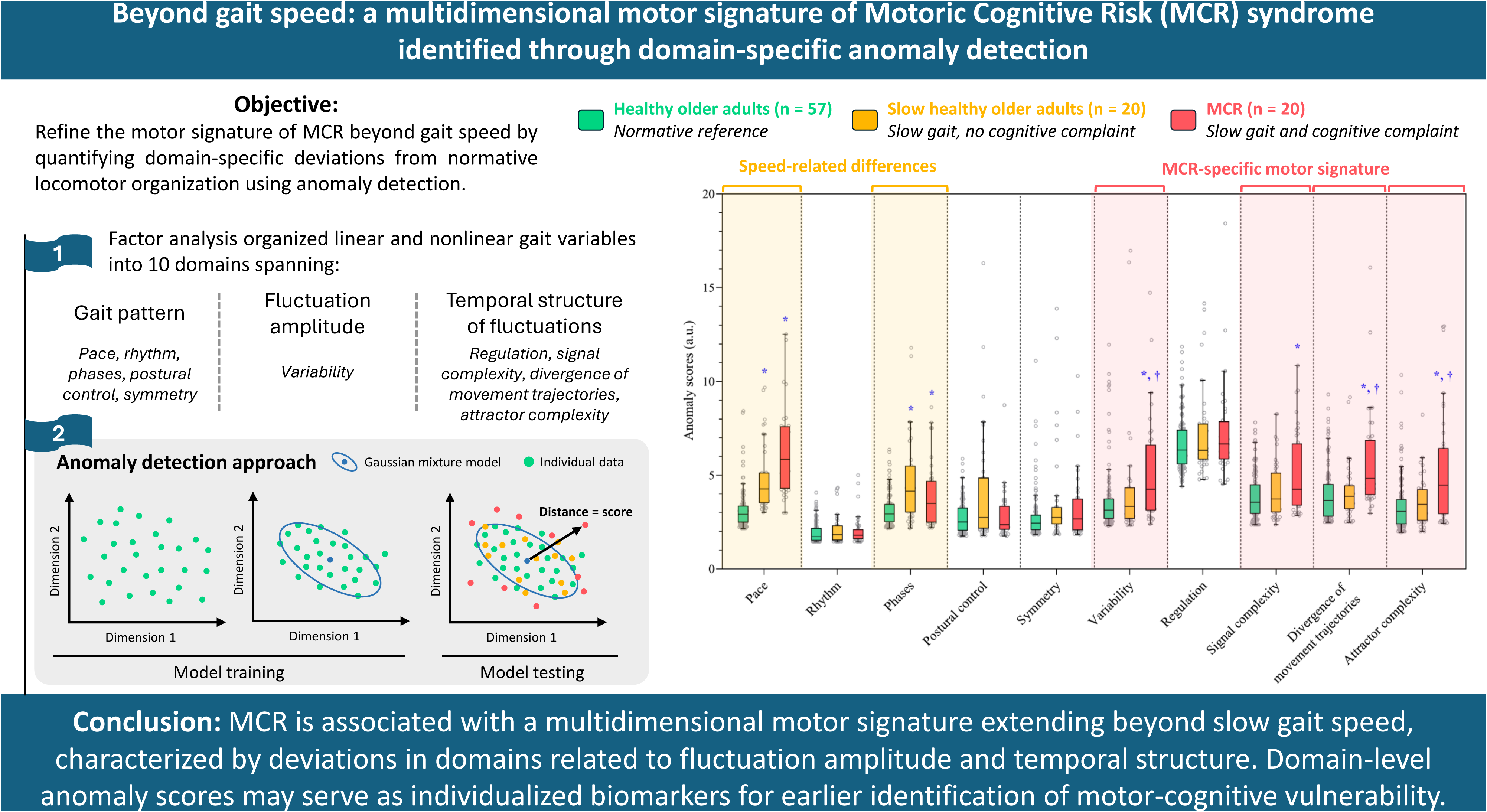

## INTRODUCTION

The demographic shift toward an aging society has led to an increase in age-related chronic conditions, including major neurocognitive disorders (NCDs). According to the World Health Organization, the global prevalence of major NCDs is currently estimated at over 55 million cases, and projections indicate that this number will rise to 78 million by the year 2030 [1]. However, subjective cognitive complaints are more frequently reported than diagnoses of major NCDs with up to 25% of adults aged 60 years and older reporting such complaints [2], compared with only 5-8% meeting diagnostic criteria for major NCDs [3]. A 14-year longitudinal study of more than 5,600 cognitively normal adults aged 50 years and older found that persistent subjective cognitive complaints were associated with a 2.2-fold increased risk of progression to mild cognitive impairment or major NCDs compared with adults without such complaints [4]. These findings suggest the existence of an intermediate population exhibiting early cognitive vulnerability despite not meeting the diagnostic criteria for major NCD.

Motoric Cognitive Risk (MCR) syndrome is one such intermediate preclinical state, defined in community-dwelling older adults aged 60 years and older by the co-occurrence of subjective cognitive complaints and a gait speed at least one standard deviation below age- and sex-adjusted reference values [5]. A meta-analysis of international cohorts estimated the prevalence of MCR in older adults to range between 6% and 10% and demonstrated that the concomitant presence of subjective cognitive complaints and slow gait speed was a more reliable predictor of conversion to major NCDs when considered together than separately [6,7].

However, gait is a complex behavior that emerges from the dynamic coordination of neuromusculoskeletal and cognitive subsystems [8–10]. Consequently, walking speed is intrinsically influenced by multiple interacting factors. Reduced speed may reflect a wide range of factors, including joint pain [11], muscle weakness [12], psychological conditions [12], sensory loss [13], inflammatory activity [14], fear of falling [15], or alterations in central motor control [16]. Accordingly, although the MCR construct has demonstrated prognostic value, reliance on gait speed as the sole motor criterion remains mechanistically nonspecific and fails to distinguish between peripheral, central, and compensatory sources of walking impairment. As a result, current MCR criteria provide only a coarse characterization of motor vulnerability and may overlook additional dimensions of gait, including gait pattern, fluctuation amplitude, and their temporal structure [7,17]. Moving beyond speed toward a more comprehensive motor signature is therefore essential to improve mechanistic interpretability, refine the identification of vulnerability profiles, and establish quantitative markers for longitudinal monitoring and evaluation of responses to preventive or therapeutic interventions.

A useful conceptual framework to characterize the motor signature is to examine gait across complementary levels of analysis that capture distinct aspects of locomotor organization and control. The first level concerns the *gait pattern*, defined by mean spatiotemporal parameters and reflecting the overall walking strategy [18,19]. The second level concerns the *amplitude of fluctuations*, defined by the variability of spatiotemporal parameters, typically quantified using linear measures of dispersion such as the standard deviation, and reflecting the consistency with which the gait pattern is reproduced from step to step [20,21]. While these two levels describe the adopted walking strategy and its temporal consistency, they provide limited insight into how gait is actively regulated, that is, the step-to-step control processes through which the neuromusculoskeletal system maintains functional walking by continuously detecting and correcting small perturbations [22]. The third level of analysis therefore concerns the *temporal structure of fluctuations*, characterized using nonlinear methods and reflecting how step-to-step fluctuations are organized across time scales.

Despite the conceptual relevance of this multilevel motor framework, most studies on MCR have focused primarily on gait pattern measures, with limited attention to fluctuation amplitude, and no studies to date have applied nonlinear methods to investigate the temporal structure of these fluctuations [17,23]. However, deriving a meaningful motor signature from these three levels of analysis remains challenging, as measures of gait pattern, fluctuation amplitude, and temporal structure are often interdependent. A single underlying process may simultaneously influence mean spatiotemporal behavior and the fluctuations around it, both in terms of their magnitude and temporal organization. Consequently, approaches based on isolated univariate comparisons may fail to capture alterations that are primarily expressed through the covariance structure among variables, leading to an incomplete characterization of gait [24]. Multivariate approaches provide a way to address this limitation by leveraging shared variance across variables and capturing latent dimensions of locomotor organization that are not apparent when variables are considered separately. A common strategy relies on factor analysis, which has consistently identified independent domains of locomotor organization, including pace, rhythm, symmetry, and postural control, with more recent work extending to stability and complexity [25–27]. Building on this domain-based representation, an important next step is to move from describing locomotor organization to quantifying individual deviations.

Machine learning methods, such as data-driven anomaly-detection models, provide a natural bridge. When trained on a healthy reference cohort, these models can learn the multivariate organization of gait within each domain and generate subject-specific deviation scores relative to that normative space [28,29]. This approach is particularly relevant for MCR, as the syndrome is heterogeneous and often expressed through subtle, distributed alterations that may not be captured by single gait metrics. Rather than testing whether a given variable differs on average between groups, deviation scoring evaluates whether an individual’s multivariate gait profile diverges from the expected domain-specific organization, thereby revealing distributed alterations that would remain undetected in conventional group-level comparisons.

The present study aims to refine the motor signature of MCR, beyond gait speed considered in isolation, by quantifying individual deviations from normative locomotor organization across multiple gait domains. Specifically, measures of gait pattern, fluctuation amplitude, and temporal structure were organized into functional domains using a data-driven framework, after which domain-specific anomaly models were trained on healthy older adults to quantify how each individual with MCR deviates from normative patterns. Because slow gait is inherently nonspecific, we additionally included a comparison group of older adults with slow gait but without subjective cognitive complaints, fulfilling only the motor criterion of MCR and providing a reference to distinguish deviations related to reduced speed from those specific to the syndrome. We hypothesized that individuals with MCR would exhibit greater deviations in domains indexing fluctuation amplitude and temporal structure, consistent with a multidimensional alteration of locomotor control rather than an isolated reduction in gait speed.

## METHODS

### Participants

Ninety-seven community-dwelling older adults aged 55 years and above were enrolled in this study. Eligibility criteria included age ≥ 55 years, right-handedness, preserved autonomy in daily living as assessed by the Katz [30] and Lawton [31] scales, and the ability to understand and comply with the study instructions. Exclusion criteria included conditions likely to interfere with gait performance or study participation: uncorrected visual or auditory impairments, active depression, history of major neurological (e.g., stroke with lasting effects) or psychiatric disorders, neuromuscular disease, severe obesity (body mass index > 34.9), and inability to provide informed consent. Within this cohort, 20 participants met the criteria for MCR syndrome, which required (i) the presence of a subjective cognitive complaint as assessed using a standardized questionnaire, (ii) slow gait speed, defined as a preferred walking speed at least one standard deviation below age- and sex-adjusted mean of the control group on the timed 10-meter walking test, and (iii) cognitive performance within the normal to subnormal range, defined by the Montreal Cognitive Assessment (MoCA) score ≥ 24 [32]. Among the 77 participants who did not meet the criteria for MCR, 20 were selected using a 1:1 nearest-neighbor matching procedure based on gait speed, age, sex, body mass index, and educational level relative to the MCR group, thereby constituting the slow healthy older adults (sHOA) group. The remaining 57 participants constituted the healthy older adults (HOA) group (Table 1). Participants were recruited through community advertisements, local associations for older adults, and memory clinics. Written informed consent was obtained in accordance with the Declaration of Helsinki, and the study protocol was approved by the regional ethics committee (IRB approval number: 2021-A02085-36).

**Table 1.**
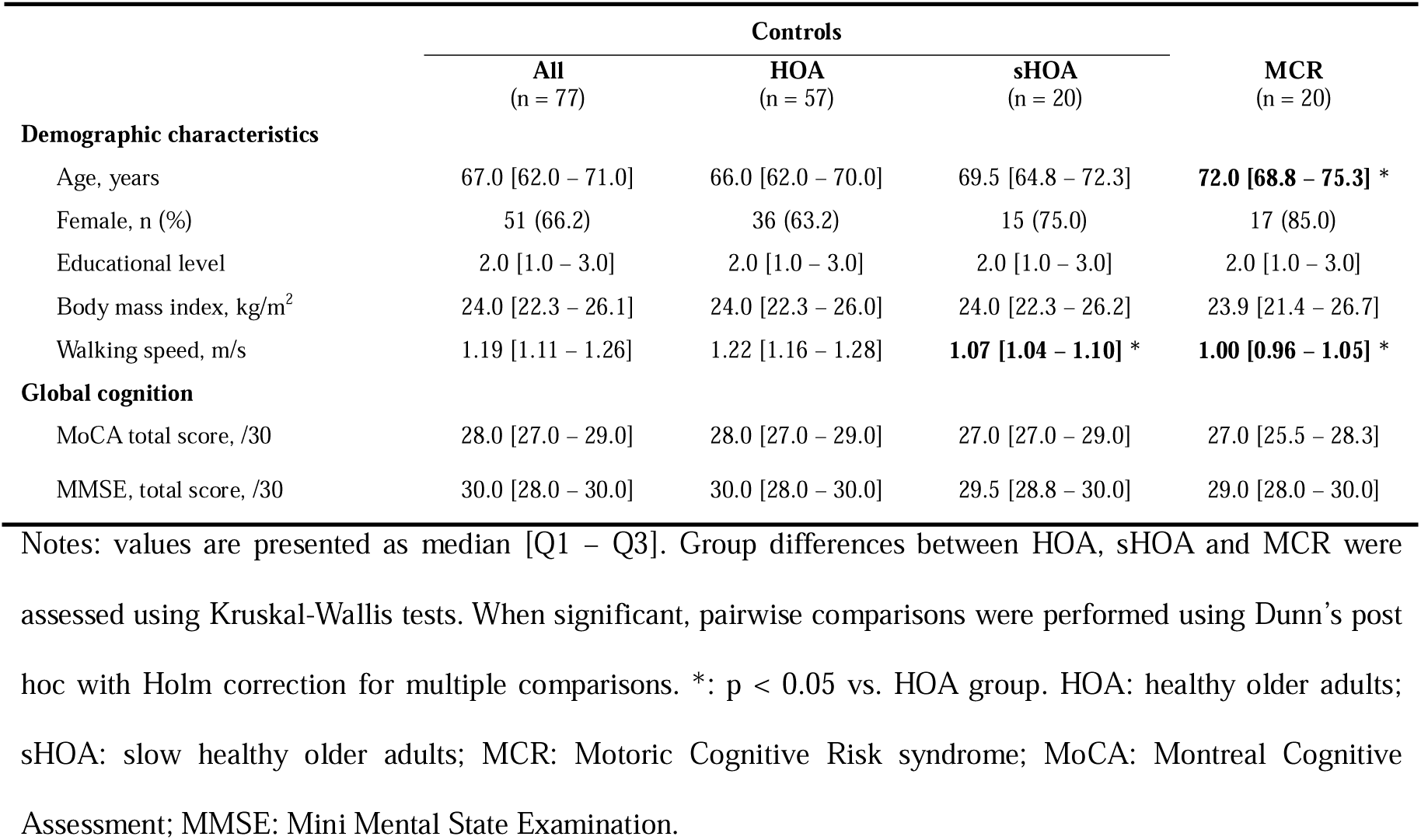
Demographic and clinical characteristics of the three study groups Notes: values are presented as median [Q1 – Q3]. Group differences between HOA, sHOA and MCR were assessed using Kruskal-Wallis tests. When significant, pairwise comparisons were performed using Dunn’s post hoc with Holm correction for multiple comparisons. *: p < 0.05 vs. HOA group. HOA: healthy older adults; sHOA: slow healthy older adults; MCR: Motoric Cognitive Risk syndrome; MoCA: Montreal Cognitive Assessment; MMSE: Mini Mental State Examination.

### Experimental procedure

Each participant completed a cognitive-locomotor protocol comprising treadmill walking under both single-task and dual-task conditions. The dual-task conditions paired walking with an adapted Stroop paradigm with three levels of difficulty [33] and will be reported in a separate publication. The present analyses focused exclusively on the two single-task walking bouts of 3 minutes 30 seconds each, which were placed at the beginning and end of the dual-task conditions. Gait assessments were conducted on an instrumented dual-belt treadmill (M-Gait, 2.0 × 0.5 m, Motekforce Link, The Netherlands) equipped with two embedded force platforms. Prior to data collection, a familiarization phase was conducted to ensure safety and comfort, and to determine preferred walking speed using the incremental protocol of Jordan et al. [34], which was then used to set the speed for all walking bouts. The treadmill was coupled with an immersive virtual-reality environment specifically developed for the research project (Fig. 1), providing controlled yet ecologically valid walking conditions. Participants were equipped with 27 reflective markers placed on anatomical landmarks according to an adapted Plug-in Gait Full Body model [35], retaining head, trunk, and lower-limb segments. Three-dimensional marker trajectories were acquired at 200 Hz using a 14-camera motion-capture system (Vero 2.2, VICON Motion Systems Ltd., Oxford, UK).

**Fig. 1.**
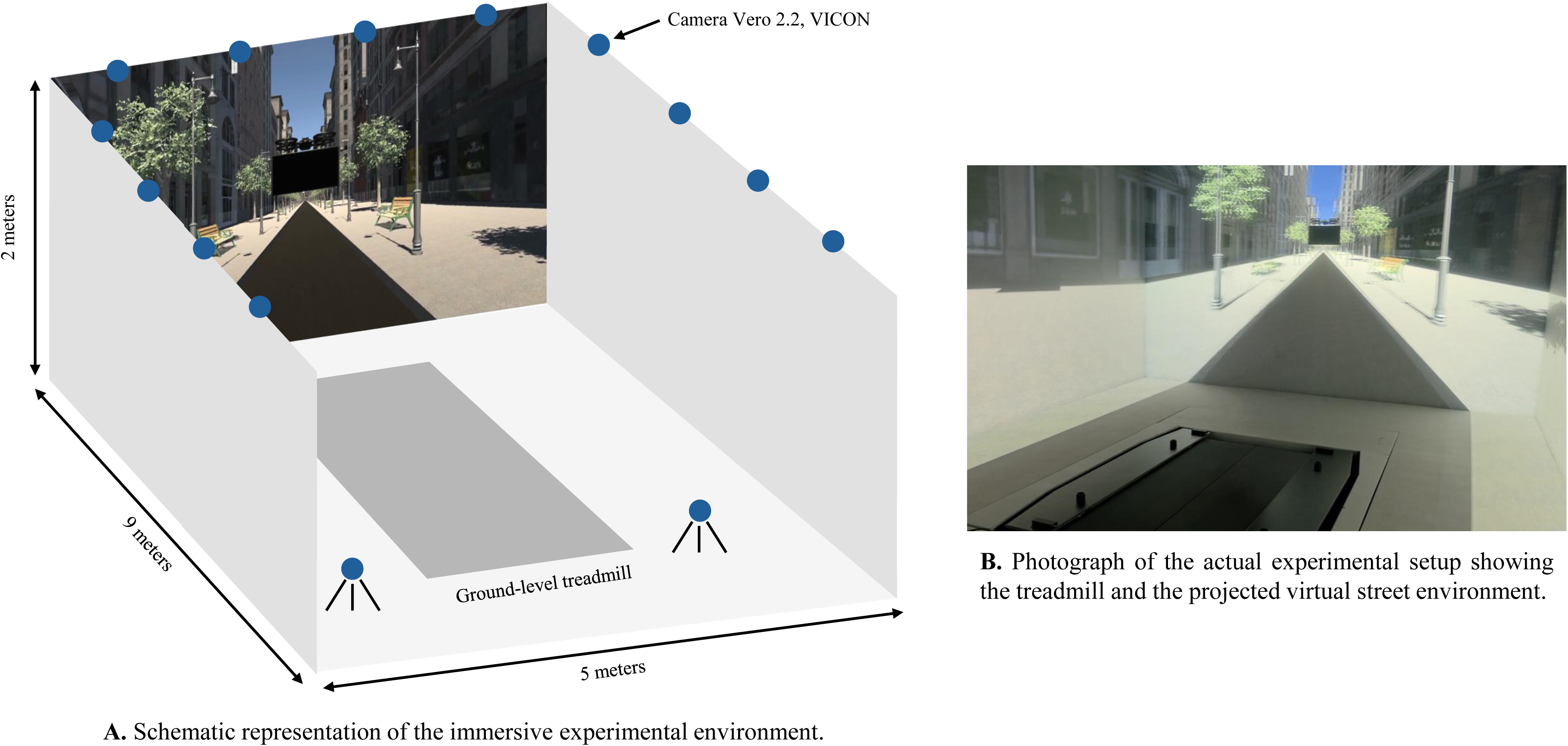
Immersive virtual-reality room and setup used for the PRESAGE study.

### Gait variables extraction

Three-dimensional marker trajectories were preprocessed to remove high-frequency noise using a fourth-order zero-lag Butterworth low-pass filter with a 10 Hz cutoff frequency [36]. Filtered trajectories were imported into a custom Python 3.12 pipeline for automated gait event detection and variable extraction. Heel-strike and toe-off events were identified from the anteroposterior heel and toe markers, enabling segmentation of each walking bout into consecutive steps. Step-by-step spatiotemporal parameters were computed from these events, including step time, step length, step width, step speed, body position, and single- and double-support durations. To ensure comparability across participants, analyses were performed on a fixed sequence of the first 284 consecutive steps after gait initiation, corresponding to the minimum number available across the cohort. Gait pattern and fluctuation amplitude were then characterized by the mean and standard deviation of each step-based time series, complemented by cadence, walk ratio, stance sub-phase percentages of single- and double-support time, and the ratio of initial-to-terminal double-support duration. The temporal structure of fluctuations was assessed using nonlinear analyses applied on step-by-step spatiotemporal and trunk acceleration time series. Specifically, long-range temporal correlations were evaluated using detrended fluctuation analysis (DFA) [37] applied to discrete step-based time series (i.e., step time, step length, step speed, step width, and body position). DFA yields a scaling exponent α, where α > 0.5 indicates persistent correlations such that deviations tend to be followed by deviations in the same direction, whereas α < 0.5 indicates antipersistent correlations such that deviations are more likely to be followed by opposite-signed deviations. In the context of treadmill walking, antipersistence is commonly interpreted as tighter step-to-step error correction of the corresponding gait variable, consistent with stronger regulation of timing and foot placement to satisfy speed and position constraints, whereas persistence reflects looser regulation that allows deviations to carry over across successive steps. In parallel, continuous signals (i.e., trunk acceleration in the anteroposterior, mediolateral, and vertical directions) were analyzed to characterize dynamic stability, regularity, and complexity. Divergence of movement trajectories was quantified using the short-term maximum Lyapunov exponent (λshort), which captures the average exponential divergence of initially neighboring trajectories in reconstructed state-space over short time scales, with larger values indicating greater sensitivity to small intrinsic perturbations and a reduced capacity to attenuate them. Primary analysis of short-term divergence was performed using the algorithm of Rosenstein et al. (1993) [38], to ensure comparability with the existing literature, and a secondary sensitivity analysis was performed using the algorithm of Wolf et al. (1985) [39], which may offer greater sensitivity for detecting subtle differences in small gait datasets [40]. Signal complexity was quantified using sample entropy (SampEn) [41], which summarizes how consistently similar short patterns in trunk accelerations signals recur through the time series within a predefined tolerance, such that lower values indicate more regular and predictable fluctuations. Finally, the Attractor Complexity Index (ACI) [42] was derived from the same data to assess long-range dynamical complexity, with higher values reflecting richer long-range temporal organization of trunk dynamics and lower values indicating a simplification of locomotor dynamics, thus complementing the other metrics. For nonlinear analyses, all preprocessing steps, state-space reconstruction parameters (i.e., embedding dimension and time delay), and algorithmic settings were optimized and held constant across participants. Full methodological details, parameter values, and computational formulas for all gait variables are provided in the Supplementary Materials.

### Domain-based structuring of gait variables

We performed a factor-analytic decomposition of the full set of gait variables, in line with prior work demonstrating that gait variables cluster into partially independent latent dimensions [26,27]. The resulting factors were then mapped onto the ten domains presented below (Table 2). This domain-level organization structured the multidimensional motor signature into functionally coherent subspaces accounting for gait pattern, fluctuation amplitude and temporal structure and formed the basis for subsequent domain-specific anomaly modeling. Full details of the factor approach and loadings are provided in the Supplementary Materials.

**Table 2.**
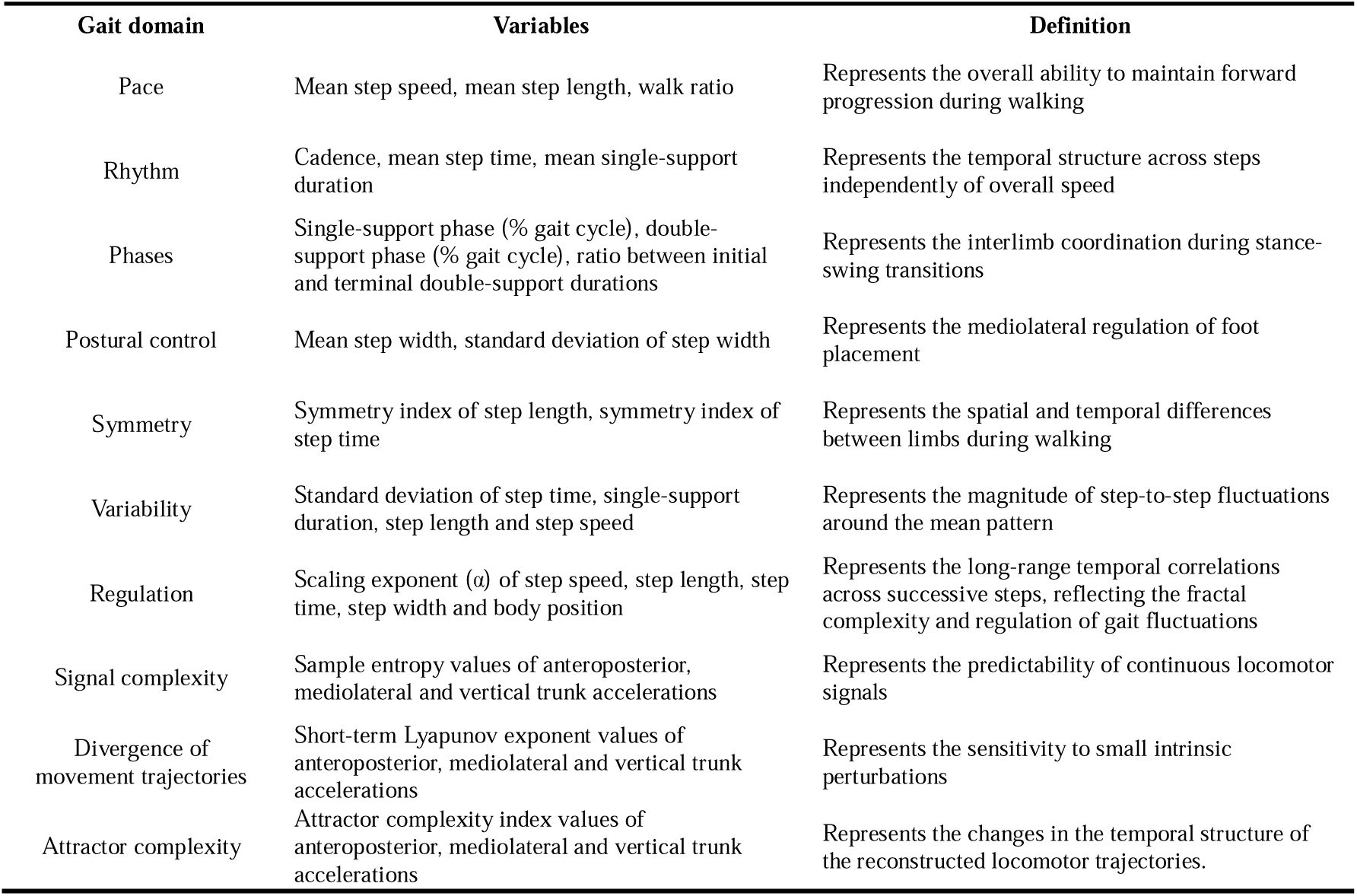
Gait domains, variables and definitions from the factor-analytic decomposition.

### Machine learning-based anomaly-detection framework

To extend beyond conventional group analysis, an unsupervised Gaussian Mixture Model (GMM) [43] anomaly-based approach was implemented to model the normative locomotor organization of the HOA group and to evaluate how and in which domain MCR and sHOA participants diverge from this reference (Fig. 2). To ensure comparability across gait domains, all domain-specific models followed the same preprocessing and training pipeline. Gait variables were first z-standardized using the HOA means and standard deviations, then projected into a lower-dimensional latent space using Principal Component Analysis (PCA) fitted on HOA only to reduce dimensionality before GMM fitting. The minimal number of orthogonal components explaining 95% of the variance in HOA was retained, and each participant’s corresponding PCA scores on these components were used as input to the GMMs (Fig. 2C). Model training and hyperparameter tuning were performed within a nested cross-validation by optimizing the median log-likelihood (i.e., computed via the GMM score_samples function in scikit-learn) across validation samples (Fig. 2D). The log-likelihood reflects how well the data fit the learned Gaussian distribution, with higher values indicating better model fit. Anomaly scores were computed for each participant and domain as the negative log-likelihood of their PCA-derived component scores under the trained GMM, higher scores indicating greater deviation from the normative control distribution (Fig. 2E). Because model parameters were estimated on HOA, these participants define the low-score reference distribution against which deviations observed in sHOA and MCR were interpreted. Both treadmill walking conditions were included in model training and evaluation to increase the number of observations and capture inter-condition variability, thereby improving the stability of variable estimates and the reliability of individual deviations. To avoid data leakage, data from a given participant was never split between training or validation samples.

**Fig. 2.**
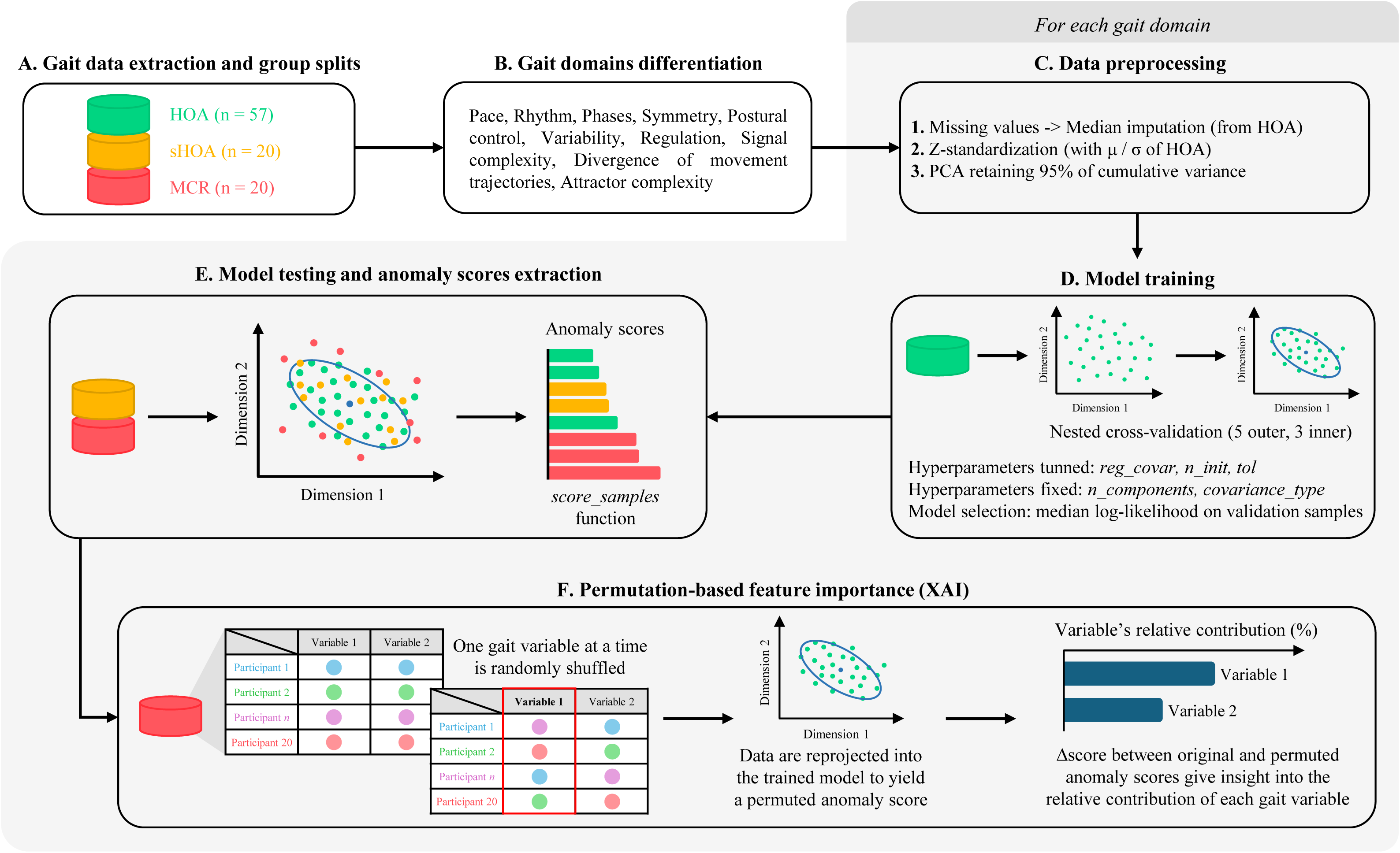
Pipeline description of the gait anomaly-detection approach.

### Explainable artificial intelligence (XAI)

A permutation-based feature-importance (PFI) analysis was conducted for each domain-specific model showing anomalies in MCR and/or sHOA [44,45] to identify which gait variables contributed most to the deviations from the normative model learned on HOA. Within each group and selected domain, gait variables were permuted one at a time using 19 complete shuffles across participants of the same group, while all other variables were kept unchanged. Anomaly scores were then recomputed using the same PCA and GMM parameters learned on HOA, thereby preserving the normative reference space. The influence of each variable was quantified as the mean absolute change in anomaly scores (i.e., Δscore) induced by the permutations. Higher values indicate that the anomaly score was more sensitive to disruption of that variable’s information, suggesting a stronger contribution to the observed deviation from the normative model. Because models were fitted separately for each gait domain, feature importance was interpreted within domains. For ranking purposes, importance values were normalized within each domain and expressed as contribution percentages (Fig. 2F).

### Statistical analyses

Gait anomaly scores were analyzed using nonparametric statistics due to non-normal distributions. Overall group effects were assessed with the Kruskal-Wallis test. When a significant omnibus effect was identified, post hoc pairwise comparisons were performed using Dunn’s test with Holm correction to control for multiple comparisons. The magnitude of pairwise group differences was quantified using Cliff’s delta (δ), a distribution-free effect size measure suitable for non-normally distributed data. Effect sizes were interpreted as negligible (|δ| < 0.147), small (0.147 ≤ |δ| < 0.33), medium (0.33 ≤ |δ| < 0.474), or large (|δ| ≥ 0.474). All statistical tests were two-tailed, and statistical significance was set at p < 0.05 after Holm correction. All analyses were performed in Python (3.12), relying on Scipy and scikit-posthocs libraries.

## RESULTS

Domain-specific gait anomaly scores differed across groups, revealing a graded profile of locomotor deviation from the normative HOA distribution (Fig. 3 and Fig. 4). In sHOA, deviations were restricted to domains closely related to gait slowing. In the *Pace* domain, anomaly scores were higher than in HOA (Kruskal-Wallis, H(2) = 90.81, p < 0.001; Dunn-Holm, p < 0.001), with a large effect size (Cliff’s δ = 0.73). Permutation analysis indicated that this deviation was driven primarily by mean step speed (52.67%), followed by mean step length (25.55%) and walk ratio (21.78%). A second, more moderate deviation was observed in the *Phases* domain (Kruskal-Wallis, H(2) = 18.11, p < 0.001; Dunn-Holm, p < 0.001), with a medium effect size (Cliff’s δ = 0.45). In this domain, contributions were mainly driven by the initial-to-terminal double-support ratio (48.75%), with smaller contributions from the single-support phase (25.92%) and double-support phase (25.33%). No other domain differed significantly between sHOA and HOA (all p > 0.05).

**Fig. 3.**
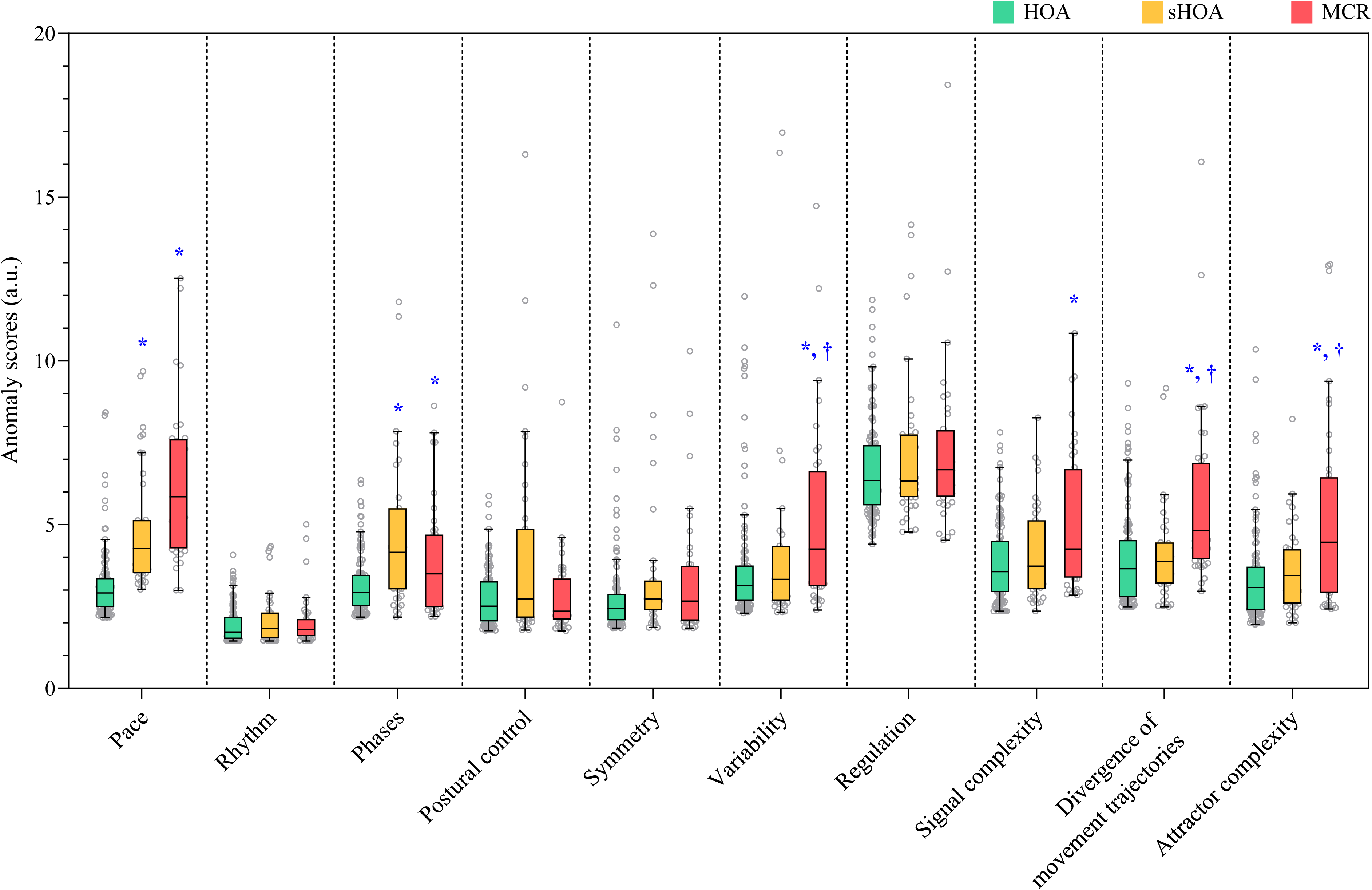
Domain-specific gait anomaly scores across groups. Anomaly scores are shown for each gait domain across the HOA (green), sHOA (yellow) and MCR (red) groups. Boxplots show median and interquartile range (Q1 – Q3); empty circle represent individual observations. *: p < 0.05 vs. HOA group; †: p < 0.05 vs. sHOA group. HOA: healthy older adults; sHOA: slow healthy older adults; MCR: Motoric Cognitive Risk syndrome.

**Fig. 4.**
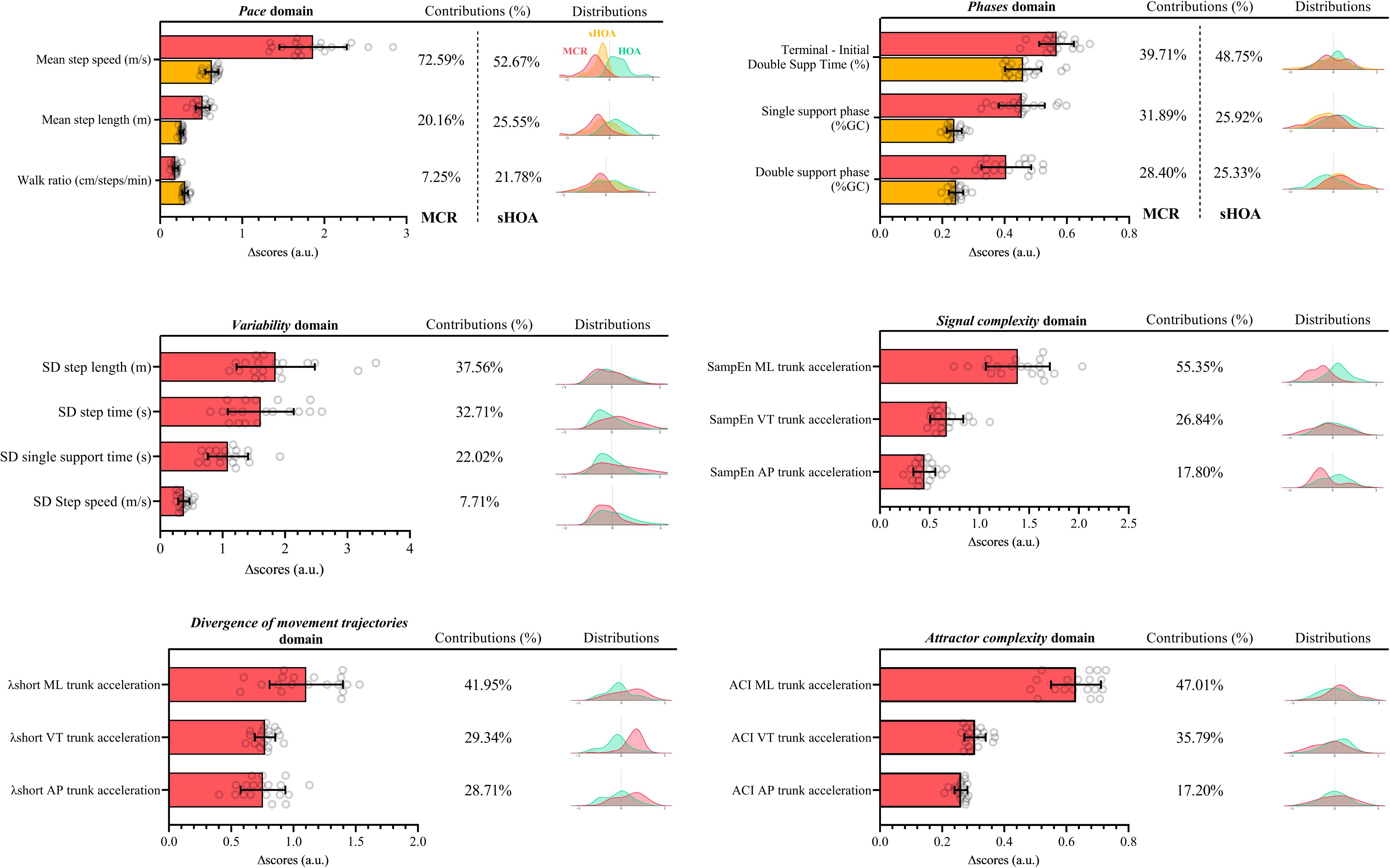
Domain-specific permutation feature importance in MCR and sHOA. Red bars and yellow bars indicate mean Δscores across 19 permutations for the MCR and sHOA groups respectively. Empty circles indicate individual permutation values. Percentages correspond to the relative contribution of each gait variable within this domain. Centered and scaled density curves are displayed for each variable to facilitate visual comparison of between-group distributional differences independently of measurement scale. sHOA: slow healthy older adults; MCR: Motoric Cognitive Risk syndrome; SD: Standard Deviation, SampEn: Sample Entropy, λshort: Short-term maximum Lyapunov exponent, ACI: Attractor Complexity Index, AP: Anteroposterior, ML: Mediolateral, VT: Vertical.

By contrast, MCR showed a broader and more specific signature of deviations from the normative HOA distribution. As in sHOA, the strongest deviation was observed in the *Pace* domain (Dunn-Holm, p < 0.001), but with a larger effect size than in sHOA (Cliff’s δ = 0.86 vs. HOA). This deviation was dominated by mean step speed (75.29%), with smaller contributions from mean step length (20.16%) and walk ratio (7.25%). A significant difference was also present in the *Phases* domain (Dunn-Holm, p = 0.048), although with a small effect size (Cliff’s δ = 0.24 vs. HOA), and contributions distributed across the initial-to-terminal double-support ratio (39.71%), single-support phase (31.89%), and double-support phase (28.40%).

Beyond these pace-related domains, MCR exhibited additional deviations not observed in sHOA. In the *Variability* domain, anomaly scores were higher in MCR than in both HOA and sHOA (Kruskal-Wallis, H(2) = 13.30, p = 0.001; Dunn-Holm, p < 0.001 and p = 0.048), with medium and small effect sizes respectively (Cliff’s δ = 0.39 vs. HOA; δ = 0.29 vs. sHOA). This deviation was mainly driven by step length variability (37.56%) and step time variability (32.71%), with smaller contributions from single-support variability (22.02%) and step speed variability (7.71%). In the *Signal complexity* domain, MCR also showed higher anomaly scores than HOA (Kruskal-Wallis, H(2) = 11.43, p = 0.003; Dunn-Holm, p = 0.002), with a medium effect size (Cliff’s δ = 0.36). Here the dominant contributor was the sample entropy of mediolateral trunk acceleration (55.35%), followed by the vertical (26.84%) and anteroposterior (17.80%) components. The largest and most specific deviations in MCR were observed in the *Divergence of movement trajectories* domain, where anomaly scores were higher in MCR than in both HOA and sHOA (Kruskal-Wallis, H(2) = 25.68, p < 0.001; Dunn-Holm, p < 0.001 for both comparisons), with large effect sizes (Cliff’s δ = 0.53 vs. HOA and 0.49 vs. sHOA). Sensitivity analyses utilizing Wolf’s algorithm yielded consistent results, confirming the group deviations observed with the Rosenstein’s algorithm (Kruskal-Wallis, H(2) = 19.29, p < 0.001, Dunn-Holm, p < 0.001 and p = 0.038 for both comparisons). The largest contributor arose from the short-term Lyapunov exponent of the mediolateral trunk acceleration (41.95%), followed by the anteroposterior (29.34%) and vertical (28.71%) components. Similarly, in the *Attractor complexity* domain, MCR showed higher anomaly scores than both HOA and sHOA (Kruskal-Wallis, H(2) = 18.48, p < 0.001, Dunn-Holm, p < 0.001 and p = 0.035), with medium effect sizes (Cliff’s δ = 0.45 vs. HOA and 0.35 vs. sHOA). This deviation was driven primarily by the attractor complexity index of the mediolateral trunk acceleration (47.01%), followed by the vertical (35.79%) and anteroposterior (17.20%) components. No other significant group effects were observed for the *Rhythm*, *Postural control*, *Symmetry* or *Regulation* domains (all p > 0.05).

## DISCUSSION

The present study shows that MCR is associated with a multidimensional motor signature that cannot be reduced to slow gait speed alone. Using domain-specific anomaly detection trained on healthy older adults, we found that individuals with MCR exhibited consistent deviations from normative locomotor organization not only in pace-related domains, but also in domains relative to fluctuation amplitude and temporal structure. While older adults with exclusively slow gait (sHOA) showed deviations primarily limited to pace, individuals with MCR displayed a unique profile characterized by domain-specific gait anomalies, namely exacerbated *Variability*, diminished *Signal complexity* and *Attractor complexity*, and greater *Divergence of movement trajectories*. As discussed below, this specific signature reflects a profound loss of locomotor adaptability and provides a mechanistic basis for the elevated risk of falls associated with this syndrome.

A major methodological strength of the present study lies in its ability to overcome a fundamental confounding factor in gait research: the pervasive influence of walking speed. Because slow gait is a defining criterion of MCR, any attempt to isolate its true motor signature must distinguish alterations inherently caused by walking more slowly from those specifically driven by the syndrome. The combination of normative deviation scoring [28,46] with a speed-matched comparison group (sHOA) offers an approach to addressing this issue. The observation that both MCR and sHOA groups exhibited higher anomaly scores in the *Pace* and *Phases* domains suggests that these domains are strongly influenced by the mechanics of slower walking itself, in line with previous findings on the effects of gait speed on mean spatiotemporal organization and gait-cycle timing [18,47]. In contrast, the additional deviations observed only in MCR across the *Variability*, *Signal complexity*, *Divergence of movement trajectories*, and *Attractor complexity* domains strongly suggest a higher-level dysregulation of locomotor control that is unlikely to be fully accounted for by reduced walking speed alone.

The deviation observed in the *Variability* domain is particularly informative because it points to altered step-to-step consistency. This domain was driven primarily by the variability of step length and step time, whereas the variability of step speed contributed less strongly. Under treadmill conditions, where progression speed is externally constrained, this pattern suggests that the alteration lies less in maintaining the overall speed output than in the fine-grained regulation of the underlying spatiotemporal stepping pattern. In practical terms, individuals with MCR appeared able to successfully maintain the imposed speed, but accomplished this at the cost of a less stable stepping consistency. This aligns with optimal control models of treadmill walking based on the Minimum Intervention Principle, which demonstrate that the nervous system exploits motor redundancy by tightly correcting deviations that alter step speed while allowing goal-equivalent fluctuations in step length and step time to persist [48]. Moreover, the strict spatial and temporal boundaries imposed by the treadmill act as a stress test that magnifies underlying syndrome-specific differences in step-to-step fluctuations. In the context of aging and cognitive vulnerability, increased fluctuation amplitude is commonly interpreted as a hallmark of diminished gait automaticity, whereby the production of a consistent stepping pattern shifts from largely automatic motor programs toward more attention-demanding control [10,49]. For individuals with MCR, who already experience cognitive vulnerability, this compensatory shift creates an imbalance between executive supply and motor demand [16]. This inability to meet the heightened cognitive requirements of walking leaves the locomotor system unable to effectively attenuate small intrinsic perturbations, likely resulting in the exacerbated step-to-step spatiotemporal variability observed in our cohort. This interpretation is clinically relevant, as greater gait variability has been consistently associated with instability, cognitive decline, and future adverse outcomes in older adults, including incident major neurocognitive disorders and falls [50–53].

Whereas the *Variability* domain captured the magnitude of step-to-step fluctuations, the trunk acceleration-derived dynamics domains provide a complementary line of evidence indicating that MCR also affects the temporal structure of these fluctuations and the control processes that shape them [22]. In the *Divergence of movement trajectories* domain, higher short-term Lyapunov exponents indicated greater sensitivity to small perturbations and a reduced capacity to attenuate intrinsic disturbances during walking [54–57]. In the *Signal complexity* domain, lower Sample Entropy values indicated that trunk acceleration fluctuations were more predictable and less complex, suggesting reduced flexibility of the underlying locomotor system rather than more efficient control [41]. In the *Attractor complexity* domain, reduced indices indicated changes in the temporal structure of the reconstructed locomotor trajectories, reflecting a simplified long-range divergence of trunk dynamics within the state space [42,58]. Taken together, these findings describe a locomotor system suffering from a dual degradation: it is globally more constrained in its dynamical repertoire, as reflected by reduced signal and attractor complexity, yet locally less able to prevent the divergence of movement trajectories, as indicated by greater short-term divergence rates. This combination suggests that the narrowing of the available motor solutions in MCR is accompanied by a failure of the fine-grained control mechanisms that normally ensure stability within the remaining operational range. This provides a striking illustration of the theoretical model of optimal movement variability [22]. While this model posits that healthy functional movement is characterized by an optimal chaotic structure and that pathology drives the system toward states that are either overly rigid or unstable, our findings reveal that MCR embodies both extremes simultaneously: a rigidification of the global locomotor pattern coupled with local step-to-step instability. This aligns with the loss-of-complexity framework in which aging and pathology progressively narrow the range of available motor solutions, reducing the system’s capacity to flexibly adapt to ongoing perturbations [59]. As recently highlighted by Mangalam (2025) [60], a rich temporal structure of variability is the true signature of adaptive potential. Thus, the diminished signal complexity and the increased divergence of movement trajectory observed in our MCR cohort reflect a profound loss of this adaptive potential, rendering the locomotor system less capable of flexible exploration and more susceptible to intrinsic and extrinsic perturbations.

Although the present study does not include neuroimaging data, the literature strongly suggests that this loss of locomotor adaptability stems from underlying structural and vascular brain abnormalities. Specifically, the MCR syndrome has been consistently associated with reduced cortical thickness in frontoparietal regions [61] and a higher burden of white matter hyperintensities [62]. These regions and white matter tracts are critical components of the neural networks responsible for the executive control of locomotion and attention [10]. Their degradation disrupts the dynamic coordination of cognitive and motor subsystems, rendering individuals with MCR particularly vulnerable to cognitive-motor interference. Consequently, under conditions that impose strict spatial and temporal boundaries (i.e., maintaining a constant speed within a confined space) and require continuous active regulation of stepping, such as treadmill walking [48], individuals with MCR may lack the executive resources necessary to efficiently monitor and correct step-to-step errors. This specific deficit in active sensorimotor regulation provides a mechanistic explanation for the domain-specific anomalies, namely the exacerbated *Variability* and the greater *Divergence of movement trajectories*, observed in our MCR cohort.

A further notable feature of these temporal structure-related results is their convergence on mediolateral trunk components. In the permutation feature importance analyses, mediolateral acceleration contributed most strongly within the *Signal complexity*, *Divergence of movement trajectories*, and *Attractor complexity* domains. This finding is clinically meaningful because mediolateral regulation during walking depends strongly on active balance control and is closely tied to the maintenance of dynamic equilibrium [54,63]. In contrast to anteroposterior progression, which is more directly constrained by treadmill mechanics and overall walking speed, mediolateral control may be more sensitive to subtle deficits in balance regulation and locomotor supervision. The predominance of mediolateral contributions therefore suggests that an important component of the MCR motor signature may involve altered balance-related gait control.

From a clinical perspective, the combined deviations in fluctuation amplitude and temporal structure define a motor profile compatible with increased vulnerability to instability and falls. Increased gait variability, greater divergence of movement trajectories, and diminished trunk signal complexity have each been associated with fall risk in older adults [54,57,64–67]. Because these alterations were specific to MCR in our cohort and were not observed in slow-gait peers without subjective cognitive complaints, they may help explain why MCR has been associated with recurrent falls and post-fall fractures more strongly than slow gait or subjective cognitive complaints considered separately [68,69].

Several limitations should nevertheless be acknowledged. First, the MCR sample size was modest, which limits statistical power to explore potential sex or age effects within the syndrome, and calls for replication in larger, multicenter cohorts. Second, although the anomaly-detection framework was well suited to the present objective, the Gaussian mixture approach still imposes assumptions on the structure of normative locomotor organization. Future work should determine whether more flexible models can capture additional heterogeneity in aging gait. Third, gait was assessed under controlled treadmill conditions. Although this limits direct generalization to overground walking, treadmill walking assessment provided important advantages by standardizing locomotor conditions and enabling the collection of long continuous recordings, which are required for a robust estimation of the temporal structure of gait fluctuations [20,38,70,71]. Furthermore, constrained conditions such as treadmill walking may act as a “stress test” that magnifies underlying subclinical and syndrome-specific differences in gait control compared to free overground walking. By taxing these specific control mechanisms, the treadmill may have facilitated the unmasking of the MCR motor signature observed in our cohort. Fourth, the cross-sectional design precludes any conclusion regarding the prognostic value of the observed gait deviations or their temporal relationship with cognitive decline. Longitudinal studies would be especially important to establish whether these domain-specific deviations provide incremental predictive value over simple gait speed alone for incident cognitive decline, falls, or progression toward major neurocognitive disorders.

## CONCLUSION

In summary, the present findings suggest that MCR is characterized by a multidimensional motor signature that extends far beyond slow gait speed alone. In addition to pace-related deviations, individuals with MCR exhibit altered fluctuation amplitude and temporal structure relative to both healthy older adults and slow-gait peers without subjective cognitive complaints. Specifically, this syndrome is characterized by increased amplitude of step-to-step variability, diminished complexity, and greater divergence of movement trajectories. These findings support the view that MCR may reflect a reorganization of locomotor control under constrained resources, resulting in reduced adaptability, particularly in the mediolateral plane, beyond gait slowing. Consequently, combining explainable anomaly detection with domain-specific normative gait modeling offers a powerful approach to refine motor phenotyping and improve risk stratification in cognitively vulnerable older adults. Integrating these domain-level deviation scores with neuropsychological, neuroimaging, and biological markers in longitudinal designs holds great promise for the earlier and more individualized detection of older adults at high risk of progressing to major neurocognitive disorders and falls.

## ACKNOWLEDGMENTS

The authors gratefully thank all study participants for their time, commitment, and invaluable contribution to this research. They also sincerely acknowledge all institutional and clinical partners whose collaboration, logistical support, and expertise made this study possible.

The authors particularly acknowledge the Gérontopôle Seine-Estuaire Normandie, the CARSAT Normandie, and the AGIRC-ARRCO Normandie for their key role in disseminating information about the study and facilitating participant recruitment. The authors warmly thank Bernard Cheru and the teams of AGIRC-ARRCO Normandie for their particularly strong involvement in identifying and recruiting potentially eligible participants.

The authors also acknowledge the COMETE laboratory (INSERM U1075, Université de Caen Normandie), the CIREVE (Centre Interdisciplinaire de Réalité Virtuelle), and the company a-gO for their essential logistical, administrative, and technical support in facilitating the implementation of the study. The authors would like to particularly thank Sophie Madeleine, director of the CIREVE, for her key role, together with Leslie M. Decker, in securing funding and acquiring the equipment necessary for the deployment of the experimental setup.

The authors also warmly thank Audrey Sultan for her key role in the clinical coordination of the PRESAGE study, including participant follow-up and her contribution to ensuring compliance with regulatory and ethical standards.

The authors also warmly thank Hervé Normand for his valuable support in participant inclusions and clinical evaluations.

## AUTHOR CONTRIBUTIONS

Leslie M. Decker (LMD) conceived and led the PRESAGE study, within which the present work is embedded, and defined the overarching research questions and experimental paradigms. LMD, Fabien Cignetti (FC), and Baptiste Perthuy (BP) jointly formulated the study objectives and hypotheses.

LMD, BP, Clément Brifault (CB), and Hugues Vinzant (HV), together with Fabien Cignetti (FC), contributed to the development of the methodological framework, including the design of the domain-based gait analysis and anomaly-detection approach and the experimental setup. Nicolas Lefèvre (NL) contributed to the development and maintenance of the immersive virtual reality environment and the associated hardware and software systems supporting motion capture and experimental implementation.

Data collection was supervised by LMD and carried out by BP, Vincent Cabibel (VC), and Rémi Laillier (RL) in the immersive platform of the CIREVE. Clinical and neurological assessments and participant eligibility were conducted by Pierre Denise (PD), whose role as physician investigator was instrumental in enabling participant inclusion in the study.

Data management and curation, including data integration, documentation, and the implementation of data preservation strategies to ensure that data remain findable, accessible, interoperable, and reusable, were supervised by LMD. VC, RL, and BP. BP performed data processing, statistical analyses, and implemented the machine learning models under the technical supervision of HV and CB.

LMD secured funding for the PRESAGE study, leading the identification of funding opportunities, the development and coordination of grant proposals, the preparation of reports to funding agencies, and overall budget structuring. LMD, together with BP, HV, and Alexandre Dalibot (AD), contributed to the preparation of the CIFRE PhD funding application, as well as to facilitating the CIFRE framework, supporting the implementation of the project, and contributing to the scientific supervision of the CIFRE PhD. FC contributed to grant writing.

LMD supervised and coordinated the overall project, including scientific direction and methodological oversight. FC, HV, and CB provided critical methodological supervision. Nick Stergiou (NS) provided critical feedback on the scientific content and contributed to conceptual refinement.

BP drafted the original version of the manuscript and prepared the tables, figures, and supplementary material. LMD and FC substantially revised the manuscript. All authors contributed to the manuscript according to their respective areas of expertise and approved the final version.

## FUNDINGS

The PRESAGE study was funded by the Caisse Régionale d’Assurance Retraite et de la Santé au Travail (CARSAT) Normandie and by grants from the Fondation Caen Normandie Santé (Crédit Agricole Normandie and Normandie-Seine, Harmonie Mutuelle, SAMMed). The project was co-funded by the European Union and the Région Normandie within the framework of the FEDER 2014–2020 Operational Programme.

This work was also supported by a CIFRE PhD fellowship (ANRT) awarded to Baptiste Perthuy, in partnership between the COMETE laboratory (INSERM U1075, INSERM / Université de Caen Normandie) and the company a-gO, co-founded by Alexandre Dalibot and Hugues Vinzant.

The study was promoted by the CHU de Caen Normandie.

## STATEMENTS AND DECLARATIONS

None

## Notes

### Competing Interest Statement

The authors have declared no competing interest.

