## Supplementary Materials for "Beyond gait speed: a multidimensional motor signature of Motoric Cognitive Risk syndrome identified through domain-specific anomaly detection"

**Appendix A: Gait event detection**

Heel-strike and toe-off events were automatically detected from the anteroposterior displacement of heel and toe markers recorded at 200 Hz. Peaks corresponding to heel-strike (local maxima in heel trajectories) and toe-off (local minima in toe trajectories) were identified using a prominence-based peak-detection algorithm implemented in Python (SciPy, *find_peaks* function). Peak detection parameters (minimum height, inter-event distance, and prominence thresholds) were optimized empirically to ensure consistency across participants and walking conditions. Detected events were visually verified for each participant to ensure the validity of the automatic detection procedure [1].

**Appendix B: Spatiotemporal gait parameters, definitions and formulas**

From the identified gait events, step-based spatiotemporal parameters were extracted to quantify the spatial and temporal organization of walking. All parameters were computed on a step-by-step basis, except for phase variables (single- and double-support phases), which were expressed as proportions of the stride cycle. For each participant, the mean and standard deviation of every parameter were calculated across 284 steps (i.e., minimum available across the cohort) to capture central tendency and fluctuation amplitude. **Table S1** summarizes the full set of step-based parameters, including their definitions and computation formulas.

**Table S1** Step-based spatiotemporal parameters, definitions and computation formulas

| **Parameters** | **Definition** | **Computation formula** |
| --- | --- | --- |
| Step time (s) | Time interval between the heel strikes of opposite feet | $Step {time}_{n}=t_{HS}^{(n+1)}-t_{HS}^{(n)}$ |
| Step length (m) | Forward distance between the heel markers of opposite feet at heel strike. | $Step {length}_{n}=\left\vert{Heel}_{AP}^{(n+1)}-{Heel}_{AP}^{(n)} \right\vert$ |
| Step width (m) | Lateral distance between the heel markers of opposite feet at heel strike | $Step {width}_{n}=\left\vert{Heel}_{ML}^{(n+1)}-{Heel}_{ML}^{(n)} \right\vert$ |
| Body position (m) | Mean lateral position of the body computed as the midpoint between left and right heel markers at each step | $Body {position}_{n}=0.5\times({Heel}_{ML}^{(n)}+{Heel}_{ML}^{(n+1)})$ |
| Step speed (m/s) | Forward velocity of each step | $Step {speed}_{n}=\frac{Step {length}_{n}}{Step {time}_{n}}$ |
| Single-support time (s) | Period when only one foot contacts the ground | $Single {support time}_{n}=t_{HS}^{(n)}-t_{TO}^{(n-1)}$ |
| Initial double-support duration (IDS, s) | Time during which both feet are on the ground immediately after heel strike, preceding contralateral toe-off | ${IDS}_{n}=t_{TO}^{(n-1)}-t_{HS}^{(n)}$ |
| Terminal double-support duration (TDS, s) | Time during which both feet are on the ground before the current foot’s toe-off, marking the end of stance | ${TDS}_{n}=t_{TO}^{(n)}-t_{HS}^{(n+1)}$ |
| Double-support time (s) | Total period within the gait cycle when both feet are in contact with the ground | $Double {support time}_{n}={IDS}_{n}+{TDS}_{n}$ |
| IDS/TDS (%) | Ratio between the durations of initial and terminal double-support phases within each stride | ${Ratio}_{IDS/TDS}^{(n)}=\frac{{IDS}_{n}}{{TDS}_{n}}\times100$ |
| Single-support phase (%) | Portion of the stride during which only one foot contacts the ground | $Single support {phase}_{n}=\frac{Single {support}_{n}}{Stride {time}_{n}}$  where: $Stride {time}_{n}=t_{HS}^{(n+2)}-t_{HS}^{(n)}$ |
| Double-support phase (%) | Portion of the stride during which both feet contact the ground | $Double support {phase}_{n}=\frac{Double {support}_{n}}{Stride {time}_{n}}$  where $Stride {time}_{n}=t_{HS}^{(n+2)}-t_{HS}^{(n)}$ |
| Symmetry index of step length and time (%) | Normalized left–right difference in mean step length and time | ${SI}_{X}=100\times\frac{\left\vert\bar{X}_{right}-\bar{X}_{left} \right\vert}{0.5\times\left\vert\bar{X}_{right}+\bar{X}_{left} \right\vert}$  where $X\in\left\{ Step length, Step time \right\}$ |
| Cadence (step/min) | Number of steps per minute | $Cadence=60 / \bar{Step time}$ |
| Walk ratio (cm/(step/min)) | Step length normalized by step frequency (cadence) | $Walk ratio= \frac{\bar{Step length}\times100}{Cadence}$ |

**Notes:** $t_{HS}^{(n)}$ and $t_{TO}^{(n)}$ denote the timestamps of the $n^{th}$ heel strike and toe-off events, respectively. ${Heel}_{AP/ML}^{(n)}$ denote the anteroposterior (AP) or mediolateral (ML) position of the heel marker at the $n^{th}$ heel-strike events. IDS = initial double support; TDS = terminal double support; SI = symmetry index.

**Appendix C: Trunk acceleration processing**

Trunk accelerations were computed from a four-marker Plug-in Gait cluster (C7, T10, CLAV, STRN) defining upper-body trunk segment. Individual marker trajectories were gap-filled by short bidirectional linear interpolation, detrended, and low-pass filtered (fourth-order zero-lag Butterworth, 20 Hz) to suppress high-frequency noise before differentiation. The filtered positions were then twice differentiated to obtain linear accelerations, followed by a 5-sample median filter to remove residual spikes. Acceleration components were expressed in the laboratory coordinate system as anteroposterior (AP), mediolateral (ML) and vertical (VT). These preprocessing choices follow established guidance for filtering optical kinematics before differentiation and for preserving step-related frequency in trunk-acceleration signals [2,3].

**Appendix D: Nonlinear analyses**

*Detrended fluctuation analysis*

Long-range temporal correlations in step-to-step fluctuations were quantified using Detrended Fluctuation Analysis (DFA). DFA was applied to step-based time series (step time, step length, step speed, step width, and body position), each containing 284 consecutive steps per participant. The signal was first integrated, then divided into non-overlapping windows of size $n$, and a second-order polynomial was fitted within each window to remove local trends. The fluctuation function $F\left( n \right)$ was computed as the root-mean-square of the residuals and evaluated across log-spaced window sizes between 16 and 96 steps, determined by DFBETA-based optimization from the initial range $n\in[4, 142]$. The scaling exponent α was then estimated as the slope of the linear regression between $\log F(n)$ and $\log n$ within this optimized range. Higher α values indicate stronger long-range correlations in step-to-step dynamics, whereas α ≈ 0.5 corresponds to uncorrelated (white-noise-like) fluctuations [4–6].

*Sample entropy*

Signal complexity was quantified using Sample Entropy (SampEn), defined as the negative logarithm of the conditional probability that two sequences of length $m$ that match within a tolerance $r$ remain similar when extended to $m+1$ points [7]. Similarity was evaluated using the Chebyshev distance, and self-matches were excluded. SampEn was computed on continuous trunk-acceleration (AP, ML, VT). The embedding dimension was fixed at $m=2$ for all signals, and the tolerance parameter was set to $r = 0.2\times SD$ of the original time series, following recommendations from previous studies [8,9]. For transparency, SampEn values were plotted across $m\in[2-5]$ and $r\in[0.10-0.50] \times SD$ to visually inspect the stability of entropy estimates and confirm that the chosen parameters fell within the plateau region typically reported in gait studies. Lower SampEn values indicate greater regularity and predictability, whereas higher values reflect increased irregularity in trunk dynamics.

*Maximum Lyapunov exponent*

Continuous trunk-acceleration (AP, ML, VT) time series were reconstructed in state-space using time-delay embedding. The time delay ($\tau$) was defined as the first minimum of the average mutual information [10], and the embedding dimension ($m$) as the smallest dimension for which the proportion of false nearest neighbors fell below 5% [11]. Both parameters were individualized for each participant and signal, following established methodological recommendations [12,13]. Each time series was then normalized to an average of 100 samples per stride as recommended by Raffalt et al. (2019) [13], ensuring consistent temporal resolution across participants and minimizing the influence of stride-to-stride duration variability.

Divergence of movement trajectories was quantified using the short-term maximum Lyapunov Exponent (λshort), which estimates the average divergence between initially neighboring trajectories in reconstructed state space. Primary estimation relied on the algorithm of Rosenstein et al. (1993) [14] to ensure comparability with the existing gait literature. For each embedded time series, the nearest neighbors were identified using the Euclidean distance while applying a Theiler window equivalent to one stride, thereby excluding temporally adjacent points from the same trajectory segment. The logarithmic divergence of these neighbors’ trajectories was computed over time, and the mean divergence curve was obtained by averaging across all reference vectors. λshort was then estimated as the slope of the linear region of this mean divergence curve, corresponding to the first 0 - 0.5 stride cycles. Higher λshort values reflect faster local divergence of trajectories and thus lower dynamic stability, whereas smaller values indicate more stable locomotor dynamics.

As a sensitivity analysis, λshort was additionally estimated using the algorithm of Wolf et al. (1985) [15], which has been reported to offer greater sensitivity for detecting subtle differences in small gait datasets [16]. In contrast to Rosenstein’s approach, which averages the logarithmic divergence across all nearest-neighbor pairs before fitting a linear regression, Wolf's algorithm follows the divergence of a single pair of neighboring trajectories over time and periodically replaces the neighbor when its separation from the reference trajectory exceeds a predefined distance threshold or when its orientation deviates by more than a predefined angle, thereby accumulating the local logarithmic divergence rate directly. The algorithm was applied with an evolution time of 10 samples (10% of a stride) and a maximum angular separation of 30°, with minimum and maximum distance thresholds adapted to the local density of the reconstructed attractor to ensure valid neighbor replacement. The same embedding parameters (time delay and embedding dimension), Theiler window, and stride-normalized time series as for the Rosenstein implementation were used to ensure comparability between the two algorithms.

*Attractor complexity index*

Long-term divergence characteristics were finally quantified using the Attractor Complexity Index (ACI) [17], derived from the same divergence curves. While the short-term Lyapunov exponent was computed over the initial 0 - 0.5 strides, the ACI was defined as the slope of the mean logarithmic divergence curve within the 4 - 10 stride intervals, capturing slower divergence dynamics of the attractor. Higher ACI values indicate a more complex gait dynamic with wider attractor boundaries allowing greater long-term divergence, whereas lower values reflect a more restricted attractor and reduced complexity of gait control [18].

**Appendix E: Exploratory factor analysis for data-driven domain definition.**

We used exploratory factor analysis (EFA) as a data-driven starting point to identify latent dimensions underlying the full set of gait variables, in line with prior factor-analytic gait models in older adults [19,20]. All candidate variables were entered simultaneously in a single EFA. Variables were summarized at the participant level, missing values were imputed using the median, and all variables were z-transformed prior to analysis. Given the exploratory intent, no pre-EFA item reduction was performed, including no redundancy-based exclusion and no Kaiser–Meyer–Olkin iterative item removal. Factor retention was determined exclusively using Horn’s parallel analysis (**Fig. S1**), retaining factors whose observed eigenvalues exceeded those obtained from random data of identical dimensions [21]. Factors were extracted using minimum residual (MINRES) estimation and rotated using an oblique promax rotation to allow correlated factors. Variables were initially assigned to the factor on which they showed the largest salient loading in the promax-rotated pattern matrix (|loading| ≥ 0.32). **Table S2** reports the promax-rotated pattern loadings together with variable communalities (h^2^), which index the proportion of variance captured by the retained factor solution.

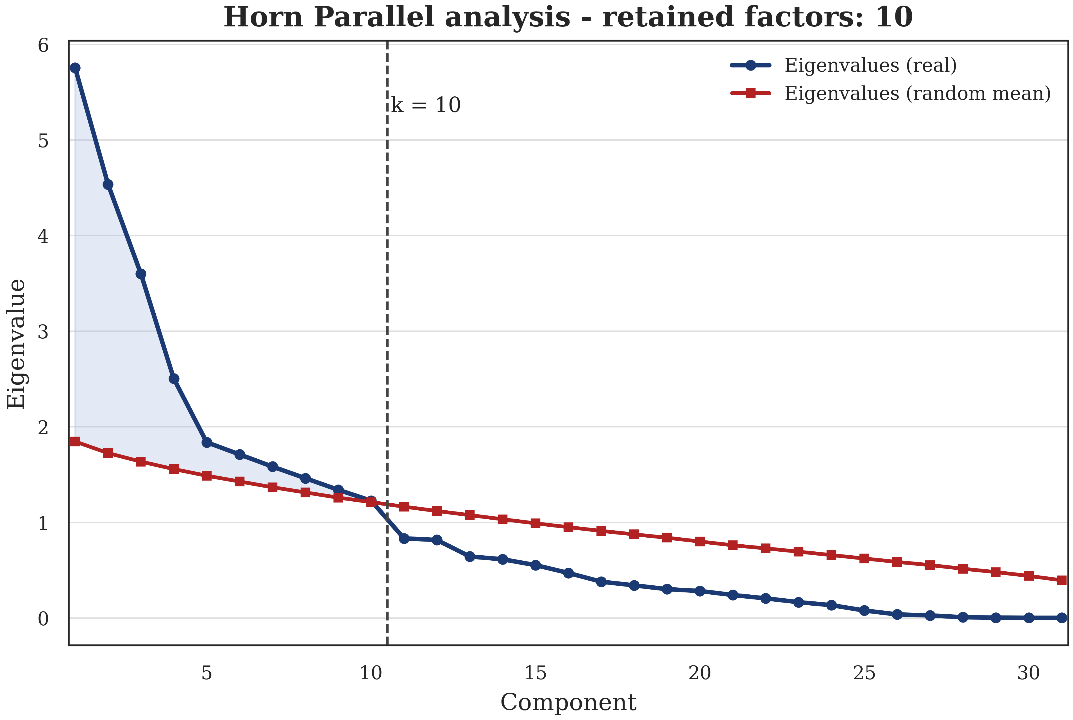

**Fig. S1** Parallel analysis for factor retention (Horn’s method). Observed eigenvalues from the empirical correlation matrix are plotted against eigenvalues obtained from random data with identical dimensions. Factors were retained when observed eigenvalues exceeded the random reference.

**Table S2** Promax-rotated pattern loadings and communalities (h^2^)

| **Variable** | **F1** | **F2** | **F3** | **F4** | **F5** | **F6** | **F7** | **F8** | **F9** | **F10** | **h^2^** |
| --- | --- | --- | --- | --- | --- | --- | --- | --- | --- | --- | --- |
| Single support time (s) | 1.09 |  |  |  |  |  |  |  |  |  | 1.30 |
| Step time (s) | 0.93 |  |  |  |  |  |  |  |  |  | 0.99 |
| Cadence (step/min) | -0.92 |  |  |  |  |  |  |  |  |  | 0.97 |
| Walk ratio (cm/steps/min) | 0.75 |  | 0.67 |  |  |  |  |  |  |  | 1.05 |
| DFA body position |  |  |  |  |  |  |  |  |  |  | 0.14 |
| SD step length (m) |  | 0.92 |  |  |  |  |  |  | 0.37 |  | 1.07 |
| SD step time (s) |  | 0.89 |  |  |  |  |  |  |  |  | 0.94 |
| SD single support time (s) |  | 0.66 |  |  |  |  |  |  |  |  | 0.57 |
| SD step speed (m/s) |  | 0.61 |  |  |  |  |  |  | 0.44 |  | 0.67 |
| Step length (m) | 0.36 |  | 0.94 |  |  |  |  |  |  |  | 1.08 |
| Step speed (m/s) |  |  | 0.88 |  | 0.34 |  |  |  |  |  | 1.02 |
| ACI trunk acceleration AP |  |  |  | 1.05 |  |  |  |  |  |  | 1.23 |
| ACI trunk acceleration V |  |  |  | 0.77 |  |  |  |  |  |  | 0.73 |
| ACI trunk acceleration ML |  |  |  | 0.65 |  |  |  |  |  |  | 0.61 |
| SampEn trunk acceleration AP |  |  |  | -0.51 |  |  |  |  |  |  | 0.47 |
| Single support phase (%GC) |  |  |  |  | 0.95 |  |  |  |  |  | 0.98 |
| Double support phase (%GC) |  |  |  |  | -0.90 |  |  |  |  |  | 0.93 |
| λshort trunk acceleration AP |  |  |  |  |  | 0.73 |  |  |  |  | 0.62 |
| λshort trunk acceleration ML |  |  |  |  |  | 0.73 |  |  |  |  | 0.67 |
| λshort trunk acceleration V |  |  |  |  |  | 0.71 |  |  |  |  | 0.60 |
| DFA step length |  |  |  |  |  |  | 1.04 |  |  |  | 1.18 |
| DFA step speed |  |  |  |  |  |  | 0.60 |  |  |  | 0.41 |
| DFA step time |  |  |  |  |  |  | 0.37 |  |  |  | 0.27 |
| Step width (m) |  |  |  |  |  |  |  | -0.77 |  |  | 0.67 |
| SampEn trunk acceleration ML |  |  |  |  |  |  |  | 0.67 |  |  | 0.60 |
| SampEn trunk acceleration V |  |  |  |  |  |  |  | 0.49 |  |  | 0.42 |
| SD Step width (m) |  |  |  |  |  |  |  | -0.32 |  |  | 0.23 |
| DFA step width |  |  |  |  |  |  |  |  |  |  | 0.14 |
| Step length symmetry index (%) |  |  |  |  |  |  |  |  | 0.76 |  | 0.69 |
| Step time symmetry index (%) |  |  |  |  |  |  |  |  |  | 0.81 | 0.77 |
| Initial to terminal double support ratio (%) |  |  |  |  |  |  |  |  |  | -0.48 | 0.38 |

**Notes:** Loadings are promax-rotated pattern coefficients. Non-salient loadings (|loading| < 0.32) are suppressed for readability. h² denotes the communality and indexes the proportion of variance in each variable accounted for by the retained factor solution. Because an oblique rotation was used, pattern coefficients may exceed 1 in absolute value. SD: Standard Deviation, DFA: Detrended Fluctuation Analysis, SampEn: Sample Entropy, λshort: Short-term maximum Lyapunov exponent, ACI: Attractor Complexity Index, AP: Anteroposterior, ML: Mediolateral, VT: Vertical.

After rotation, the promax solution yielded factors that mapped well onto established gait constructs, separating step-timing measures (cadence, mean step time, mean single-support time), step-to-step variability (standard deviations of step time, single-support time, step length, and step speed), pace (mean step speed and mean step length), and stance support phases (single- and double-support percentages with opposite loadings). Nonlinear outcomes also clustered by construct, with DFA exponents forming a regulation factor, short-term Lyapunov exponents forming a divergence of movement trajectories factor, and ACI measures forming a complexity factor. A small number of variables were reassigned a posteriori to improve functional coherence and clinical interpretability, primarily for variables showing cross-loadings and when their biomechanical meaning was more consistent with an adjacent domain. Walk ratio was grouped under Pace because it reflects step length relative to cadence and is commonly interpreted as an efficiency-related descriptor of forward progression. The ratio between initial and terminal double-support time was assigned to Phases because it captures within-stance organization rather than symmetry. SampEn measures were pooled across axes into a single signal complexity domain because they quantify the same construct of trunk-signal regularity despite direction-specific covariance. DFA step width and DFA body position were retained within the other DFA exponents to preserve a unified representation of long-range correlations in step-to-step regulation.
